# A fungal effector hijacks conserved plant GSK3-like kinases to reprogram host cell identity

**DOI:** 10.64898/2026.09.15.751668

**Authors:** Mamoona Khan, Aladár Pettkó-Szandtner, Pouria Bahrami, Hua Liu, Gregor Langen, Alga Zuccaro, Bing Yang, Armin Djamei

## Abstract

Stable developmental programs normally resist reprogramming, yet pathogens can override these constraints to induce profound tissue and cellular remodeling. How microbial pathogens access conserved developmental signaling networks to reprogram host growth and differentiation remains poorly understood. Here, we show that the maize smut fungus *Ustilago maydis* deploys the effector Iag1 to reprogram host cell fate by targeting the conserved GSK3-like kinase BIN2. Iag1 contains a PPNT short interaction motif found also in plant viral effectors that mediates its association with BIN2 and related maize GSK3-like kinases and is essential for effector activity and full fungal virulence. This interaction attenuates BIN2-dependent signaling, reduces phosphorylation of the BIN2 substrate BES1, and triggers extensive transcriptional reprogramming consistent with coordinated perturbation of brassinosteroid, auxin, and other BIN2-regulated developmental pathways. These changes result in extensive epidermal developmental reprogramming, including loss of stomatal identity, altered cell division orientation, and aberrant cell expansion. Together, our findings identify GSK3-like kinases as conserved developmental signaling hubs that pathogens can exploit to unlock host developmental plasticity and suggest that short motifs provide an evolutionarily flexible strategy for targeting these central regulatory nodes.

## Main

The maintenance of differentiated cell identity is a defining feature of multicellular organisms ^1^, yet this stability can be actively reversed in specific developmental and pathological contexts ^2–4^. While such plasticity is often attributed to endogenous regulatory programs ^2^, whether and how microbial pathogens directly reprogram host cell identity remains a fundamental open question.

Pathogen-induced gall formation is one of the most striking examples of developmental plasticity in plant-parasite interactions, in which differentiated tissues change cell proliferation, growth rate, and morphology to generate specialized organs that support pathogen colonization ^5,6^. Although these developmental alterations have long been attributed to uncontrolled proliferation, it is becoming increasingly clear that pathogens coordinate changes in cell fate, growth direction, and differentiation during gall development by exploiting innate host developmental flexibility ^6,7^. These changes are orchestrated by pathogen-derived effector proteins ^8,9^, which frequently target host regulators controlling hormone signaling, transcription, and development ^10–12^. However, the molecular mechanisms by which these effectors coordinate host developmental reprogramming remain poorly understood. The biotrophic fungus *Ustilago maydis*, the causal agent of corn smut disease, is an excellent model for studying pathogen-induced developmental reprogramming ^13–15^. During infection, *U. maydis* induces prominent galls on maize leaves and stems through the delivery of fungal effector proteins into host cells. Several effectors have been linked to immune suppression ^16–18^, metabolic reprogramming ^19,20^ and hormone signaling ^21–25^, whereas more recently identified effectors regulate host developmental programs by targeting components controlling cell proliferation and pluripotency^26–28^. Moreover, increasing evidence suggests that individual effectors promote gall formation in an organ- or cell-type-specific manner ^29–31^. Nevertheless, the molecular targets and mechanistic functions of most *U. maydis* effectors remain unknown. In particular, it remains unclear which effectors directly reprogram host cell fate and growth orientation during gall formation in maize.

Plants continuously integrate endogenous developmental programs with environmental and biotic cues to maintain growth and tissue organization. A central regulator of these processes is the glycogen synthase kinase 3 (GSK3)-like kinase BR INSENSITIVE 2 (BIN2). Initially identified as a negative regulator of BRASSINOSTEROID (BR) signaling through phosphorylation of the transcription factors BRASSINAZOLE RESISTANT 1 (BZR1) and BRI1-EMS-SUPPRESSOR 1 (BES1/BZR2), BIN2 is now recognized as a multifunctional developmental signaling hub that integrates numerous regulatory pathways ^32–34^. Beyond BR signaling, BIN2 regulates auxin responses, stomatal development, vascular differentiation, cell division, cytoskeletal organization, and multiple stress-response pathways through phosphorylation of diverse downstream substrates ^35–39^. Consequently, modulation of BIN2 activity has the potential to alter plant growth, differentiation, and tissue patterning. Its central position at the intersection of multiple developmental networks makes BIN2 an attractive target for pathogen-mediated developmental manipulation. So far, only geminiviral effectors have been reported to target GSK3-like kinases to manipulate host development ^40^.

Here, we identify and characterize a previously unstudied *U. maydis* effector, UMAG_05824 (hereafter Inducer of asymmetric growth 1, Iag1), which is specifically expressed in fungal hyphae colonizing leaf gall tissue. We show that Iag1 is secreted during biotrophic growth, contributes quantitatively to fungal virulence, and is sufficient to induce profound developmental abnormalities in the monocot host plant maize but also in the model dicot plant *Arabidopsis thaliana*. Using biochemical, genetic, transcriptomic, and cellular approaches, we demonstrate that Iag1 interacts with BIN2 and related maize GSK3-like kinases through a short PPNT interaction motif that is also present in unrelated plant viral effectors and in the metazoan scaffold protein AXIN2. Together, our findings establish BIN2 as a conserved developmental signaling hub exploited by a fungal effector to reprogram host development.

## Results

### *Iag1* is expressed during gall development and is sufficient to induce ectopic cell proliferation and disrupt epidermal patterning

To identify *U. maydis* effectors that contribute to developmental remodeling during gall formation, we examined published infection-stage transcriptomes for candidate effectors whose expression coincides with tumor-like development in the host plant maize ^30,41^. These analyses identified Iag1 as a highly expressed, previously uncharacterized candidate effector. Iag1 expression increased between two to twelve days post-infection in maize leaves infected with wild-type strains, peaking at four days post-infection, coincident with the onset of gall formation (Fig. 1a) ^41^. Consistent with this, Iag1 was among the most highly induced effector genes in gall tissue ^30^. To determine its spatial expression pattern, we generated a reporter strain expressing mCherry under the control of the native Iag1 promoter. Confocal microscopy four days post-inoculation revealed promoter activity in fungal hyphae colonizing epidermal (E), mesophyll (M), bundle sheath (BS), and vasculature, indicating that the effector is expressed broadly in hyphae colonizing maize tissue during gall formation (Fig. 1b). These observations suggested that Iag1 contributes directly to host developmental reprogramming during gall formation.

**Figure 1.**
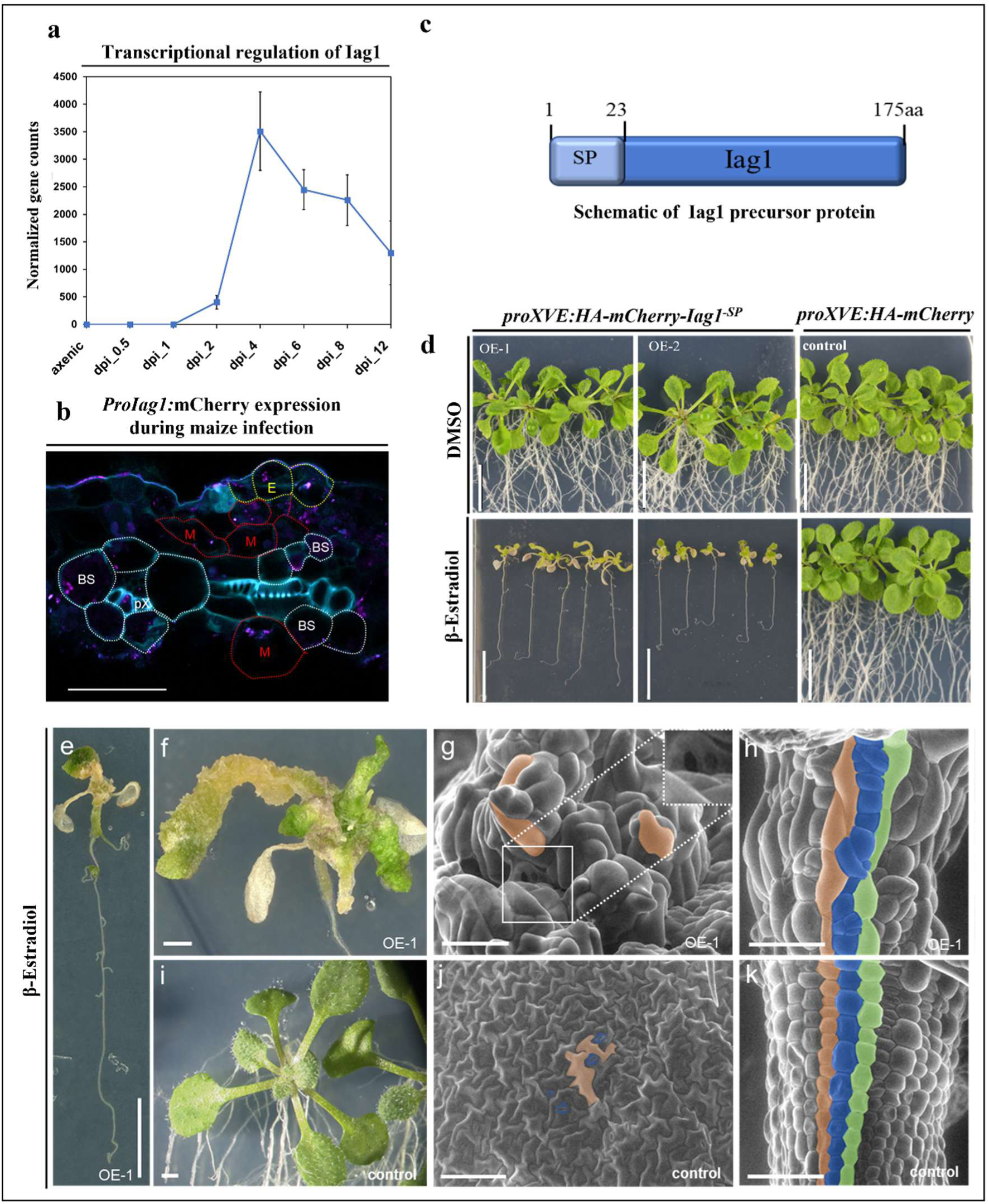
*Inducer of asymmetric growth 1 (Iag1)* is enriched in gall tissue *of U. maydis-* infected maize leaves and induces developmental reprogramming *in planta*. (a) Transcript abundance of Iag1 at different stages of *U. maydis* infection (days post-infection, dpi). Data represent three independent biological replicates; error bars indicate standard deviation (SD)^41^. **(b)** Confocal micrograph of a transverse section of a maize leaf infected with a *U. maydis* strain expressing mCherry under the control of the native Iag1 promoter. Magenta fluorescent signals identify fungal hyphae expressing the Iag1 promoter reporter, which were observed in epidermal (E), mesophyll (M), bundle sheath (BS), and protoxylem (pX)-associated tissues at 4 dpi. Scale bar, 50 µm. **(c)** Schematic representation of the Iag1 protein showing the predicted N-terminal signal peptide (SP). Numbers indicate amino acid positions. **(d)** Morphology of two independent transgenic *Arabidopsis thaliana* lines expressing intracellular Iag1 lacking the fungal signal peptide (*Iag1^-SP^*) and HA-mCherry control plants 12 days after induction with 10 µM β-estradiol. **(e,f,i)** Representative digital microscope images and **(g,h,j,k)** scanning electron micrographs of β-estradiol-induced seedlings showing ectopic epidermal cell proliferation in leaves and increased cell proliferation, cell enlargement, and tissue disorganization in hypocotyls of Iag1^-SP^-expressing plants compared with controls. Individual cells are pseudo-colored to highlight cell morphology and identity. Scale bars, 5 mm (e), 1 mm (f,i), 100 µm (g,j), and 200 µm (h,k).

Protein effectors secreted by a plant-colonizing microbe might act either in the apoplast (apoplastic effectors) or be further translocated into the plant cell (symplastic effectors). To test if Iag1 has a conserved plant target and to identify a putative apoplastic or symplastic function, we expressed in the model plant *Arabidopsis thaliana* Iag1 under a β-estradiol-inducible promoter either as an intracellular form lacking the SignalP 6.0 ^42^ predicted fungal secretion signal (*Iag1^-SP^,* Fig. 1c) or as a secreted form fused to a plant signal peptide. Induction of the secreted form did not produce detectable developmental changes (Fig. S1). In contrast, intracellular expression of *HA-mCherry-Iag1^-SP^* in multiple independent transgenic plant lines induced pronounced developmental defects, including ectopic callus-like outgrowths on leaf surfaces accompanied by severe chlorosis and shoot paling compared with control plants (Fig. 1d-f,i).

Scanning electron microscopy revealed extensive epidermal hyperproliferation in *Iag1*^-SP^-expressing plants, resulting in dense clusters of small pavement cells compared with control plants (Fig. 1g,j). Guard cells were largely absent from affected regions, with only occasional pore-like openings remaining (Fig. 1g). Similar developmental abnormalities were observed in hypocotyl tissues, which showed irregular cell divisions, enlarged cells, and disrupted tissue organization compared with controls (Fig. 1h,k). In addition, root growth was largely inhibited following Iag1 induction (Fig. 1d,e). Together, these findings demonstrate that intracellular Iag1 is sufficient to induce extensive developmental reprogramming, characterized by altered epidermal organization, enhanced cell proliferation, and disruption of normal stomatal development.

### *Ustilago maydis* disrupts epidermal patterning and growth during gall formation

Previous studies of *U. maydis*-induced gall development have focused primarily on mesophyll- and bundle sheath-derived galls, whereas the developmental response of the epidermis has remained largely unexplored ^30^. We therefore examined epidermal organization during gall formation using scanning electron microscopy and confocal imaging (Fig. 2 and S2).

**Figure 2.**
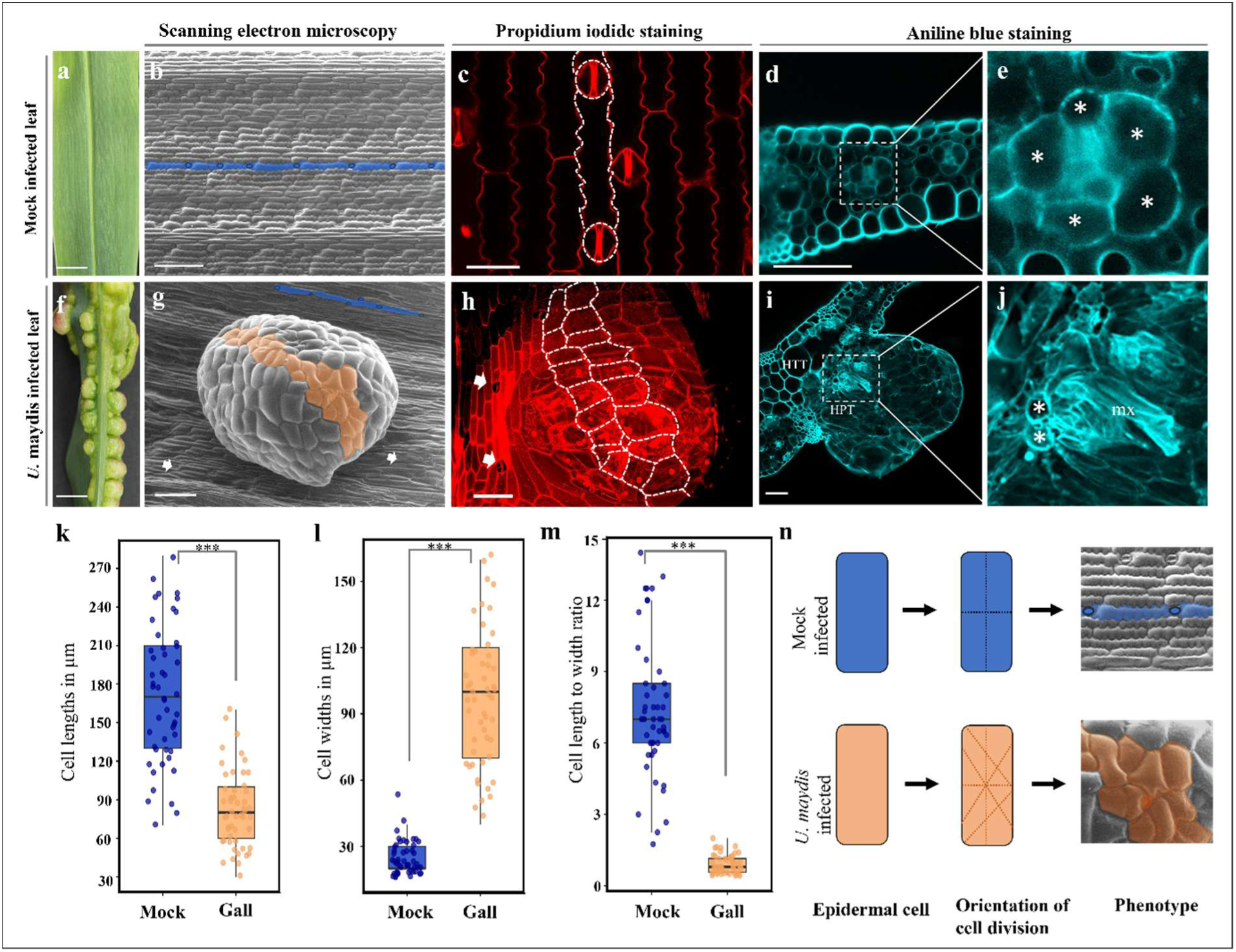
*Ustilago maydis* induces epidermal cell fate reprogramming in maize leaves. **(a-e)** Morphology of mock-inoculated maize (inbred line B73) leaves. (**f-j)** Morphology of maize leaves inoculated with the compatible *U. maydis* strains FB1 and FB2. **(b, g)** Scanning electron micrographs of the abaxial leaf epidermis 7 days post-inoculation (dpi). Blue pseudo-colored cells highlight the characteristic rectangular morphology of pavement cells in mock-treated leaves, whereas apricot pseudo-colored cells indicate the altered morphology of epidermal cells within gall tissue. Images were acquired from 14-day-old seedlings. **(c, h)** Confocal micrographs of propidium iodide-stained leaf epidermis from mock-treated (**c**) and *U. maydis-*infected leaves (**h**). White outlines indicate epidermal cell boundaries, and arrows indicate stomata adjacent to gall tissue. Panel h shows a Z-stack projection. **(d, e, i, j)** Confocal micrographs of transverse leaf sections stained with aniline blue (cyan). Asterisks indicate bundle sheath cells. HTT, mesophyll-derived hypertrophic tumor cells; HPT, bundle sheath-derived hyperplastic tumor cells; mx, metaxylem. **(k)** Quantification of pavement cell length and **(l)** width in mock-treated and gall tissues. **(m)** Length-to-width ratios of pavement cells, demonstrating a shift from anisotropic to isotropic growth following *U. maydis* infection. **(n)** Graphical representation of cell division orientation in mock versus *U. maydis-*infected maize leaf pavement cells. Scale bars, 5 mm (a,f), 200 µm (b,g), 50 µm (c), and 100 µm (d,h,i). Data in panels k–m are presented as box plots with individual data points overlaid. Statistical significance was determined using an unpaired two-tailed Student’s t-test (P < 0.0001).

In mock-infected leaves and in non-gall regions of infected leaves, epidermal cells were arranged in longitudinal files aligned with the leaf axis and displayed regularly spaced stomata separated by characteristic lobed pavement cells of relatively uniform size (Fig. 2a-d, k-m). In contrast, epidermal cells within galls lost this highly ordered organization. Cells became predominantly isodiametric, lacked the characteristic lobes of mature pavement cells, and exhibited pronounced size heterogeneity, consistent with a transition from anisotropic to isotropic growth (Fig. 2f-i).

This developmental reprogramming was accompanied by marked defects in epidermal patterning. Guard cells were largely absent from mature galls, and only occasional pore-like openings were detected compared with the surrounding non-infected epidermis (Fig. 2g-h, S2), indicating disruption of normal stomatal development. In addition, we observed previously described tracheary element-like structures at positions corresponding to former bundle sheath cells (Fig. 2d,e,i,j), suggesting ectopic differentiation events within gall tissues. Finally, whereas cell divisions in the epidermis of the mock-treated leaves were predominantly oriented perpendicular to the longitudinal growth axis, newly formed cell walls within the gall epidermis were oriented randomly, indicating a loss of coordinated division-plane control during gall development (Fig. 2k-m).

Together, these observations demonstrate that *U. maydis* infection profoundly reprograms epidermal development by disrupting cell growth, stomatal patterning, and division orientation during gall formation.

### Iag1 induces transcriptional remodeling of hormone signaling and developmental programs

The ability of intracellular Iag1 to induce extensive developmental abnormalities in a non-host dicot plant suggested that the effector targets a conserved regulatory mechanism controlling plant development. To identify host pathways perturbed by Iag1, we performed RNA sequencing (RNA-seq) on 12-day-old *Arabidopsis thaliana* seedlings expressing *HA-mCherry-Iag1^-SP^* under a β-estradiol-inducible promoter. Transgene expression was induced 4 h prior to sampling, and transcriptomes were compared with identically treated β-estradiol-inducible HA-mCherry control plants. Differential expression analysis (adjusted P < 0.05, |log₂ fold change| ≥ 1; Supplementary Dataset 1) identified 952 differentially expressed genes, including 593 upregulated and 359 downregulated transcripts (Dataset S1, Fig. S3a), demonstrating extensive transcriptional remodeling following Iag1 induction.

Gene Ontology and KEGG pathway analyses identified significant enrichment of genes associated with Brassinosteroid (BR) biosynthesis, metabolism, and signaling among differentially expressed transcripts (Fig. 3a, S3b, Dataset S2). Several core BR biosynthetic genes, including DWF4, CPD, CYP90D1, and BR6OX2, were coordinately downregulated, whereas established BR-responsive genes such as BEE3 were induced. This transcriptional signature is consistent with increased BR signaling output, which is accompanied by negative feedback repression of BR biosynthetic genes. Beyond BR-associated genes, *Iag1^-SP^*expression triggered extensive changes in additional hormone-responsive pathways. In particular, numerous auxin-responsive genes, including members of the SAUR, IAA, and GH3 families, were strongly upregulated (Fig. 3a, c; Fig. S3c; Dataset S3). Genes involved in cell cycle regulation, cell wall organization, and epidermal development were also differentially expressed, consistent with the developmental abnormalities observed in *Iag1^-SP^*-expressing plants (Fig. 3a).

**Figure 3.**
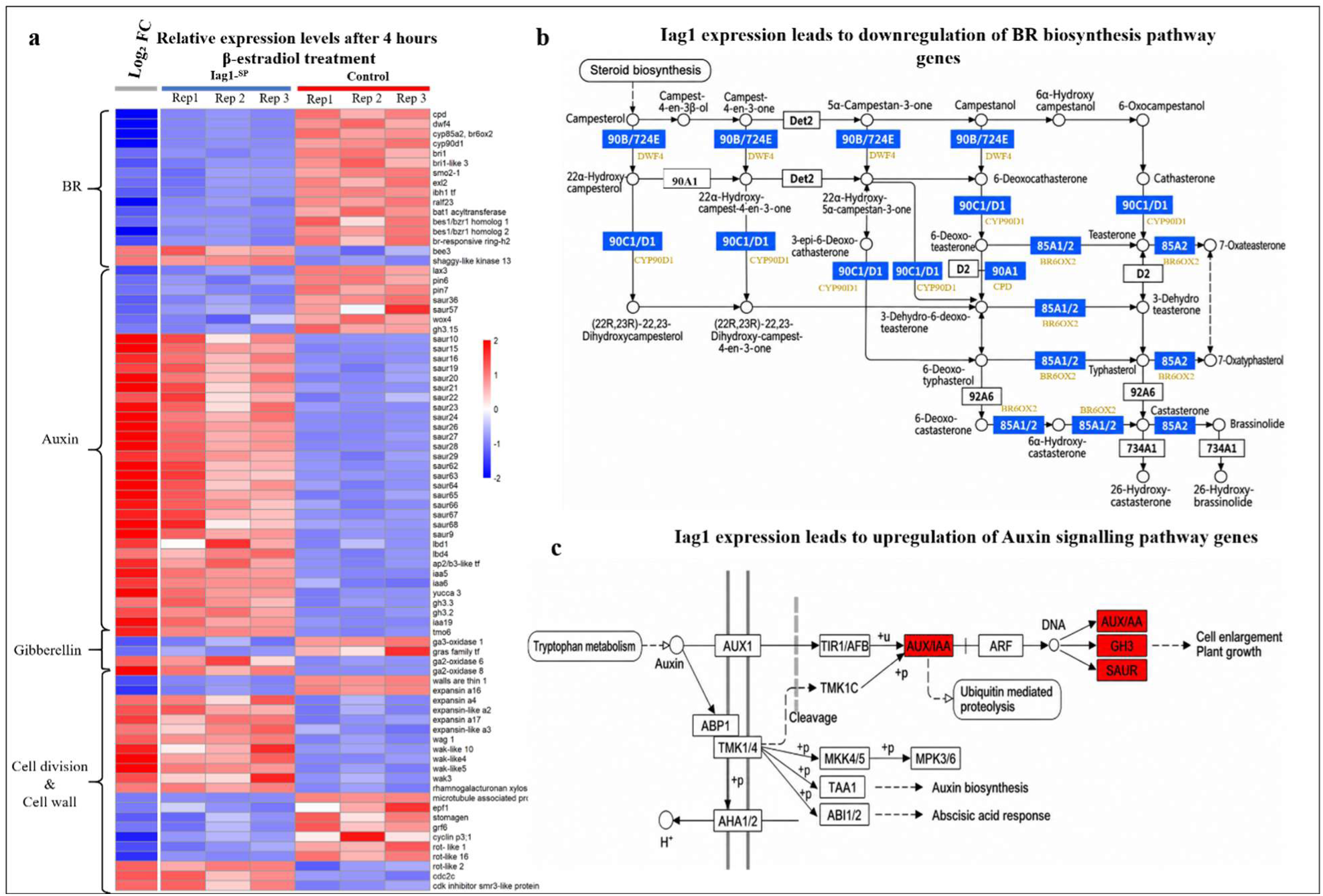
Inducer of asymmetric growth 1 (Iag1) reprograms host transcription and alters hormone biosynthesis and signaling pathways. **(a)** Heatmap showing the expression profiles of selected genes differentially expressed between β-estradiol-induced *HA-mCherry-Iag1^-SP^* and identically treated *HA-mCherry* control plants. Genes were selected based on DESeq2 differential expression analysis (adjusted P < 0.05, |log₂ fold change| ≥ 1). The leftmost column represents the DESeq2 log₂ fold change (Iag1 versus HA-mCherry), whereas the remaining columns display row Z-scores calculated from DESeq2-normalized gene counts for individual biological replicates, with Z-scores calculated separately for each gene across all samples. **(b)** KEGG pathway visualization of brassinosteroid (BR) biosynthesis generated using ShinyGO. Blue and white boxes represent enzymes in the pathway. Blue boxes indicate enzymes encoded by genes that were significantly downregulated in *Iag1^-SP^*-expressing plants compared with the control, whereas white boxes indicate enzymes whose corresponding genes were not differentially expressed. Orange labels denote the corresponding Arabidopsis gene names. **(c)** KEGG pathway visualization of auxin signaling generated using ShinyGO. Red and white boxes represent components of the auxin signaling pathway. Red boxes indicate components encoded by genes that were significantly upregulated in *Iag1^-SP^*-expressing plants compared with the control, whereas white boxes indicate components whose corresponding genes were not differentially expressed.

Together, these transcriptomic analyses demonstrate that intracellular Iag1 induces extensive developmental and hormone-responsive transcriptional remodeling. The coordinated repression of BR biosynthetic genes, activation of BR-responsive transcription, and induction of auxin-responsive gene expression suggest that Iag1 perturbs a central regulatory mechanism integrating multiple developmental signaling pathways.

### Iag1 targets conserved GSK3-like kinases through a PPNT interaction motif to attenuate GSK3-like kinases-dependent signaling

The broad developmental and transcriptional changes induced by Iag1 suggested that the effector targets a conserved regulatory component coordinating multiple developmental pathways. To identify host proteins associated with Iag1, we purified protein complexes from *A. thaliana* seedlings expressing inducible *2×Myc-Iag1^-SP^* and analyzed them by co-immunoprecipitation followed by liquid chromatography-tandem mass spectrometry (Co-IP/LC-MS). Protein complexes isolated from identically treated inducible 2×Myc-GFP plants served as negative controls (Fig. S4a). Co-IP/LC-MS identified a limited number of proteins specifically enriched with Iag1 relative to the GFP control (Fig. S4B). To determine whether these candidates directly interact with Iag1, we performed yeast two-hybrid (Y2H) assays. These assays identified the GSK3-like kinase BIN2 as a direct interactor of Iag1, whereas none of the remaining candidates showed detectable interaction (Fig. S4C).

As maize is a natural host of *U. maydis*, we next investigated the interaction of Iag1 with maize orthologs of BIN2. Y2H assays demonstrated that Iag1 interacts with the maize BIN2 homologs ZmGSK1, ZmGSK4, and ZmGSK8 (Fig. 4a,b), indicating that this interaction is conserved between non-host and native host systems. To validate these interactions *in planta*, we performed bimolecular fluorescence complementation (BiFC) assays in *Nicotiana benthamiana*. Co-expression of *Iag1^-SP^* fused to the C-terminal Venus (CVenus) fragment with BIN2 or the three maize GSK3-like kinases fused to the N-terminal Venus fragment (NVenus) resulted in strong fluorescence in epidermal cells, whereas no signal was detected with the control (Fig. 4c). Together with the Y2H results, these assays support direct interaction of Iag1 with BIN2 and related maize GSK3-like kinases and confirm that these proteins associate in plant cells.

**Figure 4.**
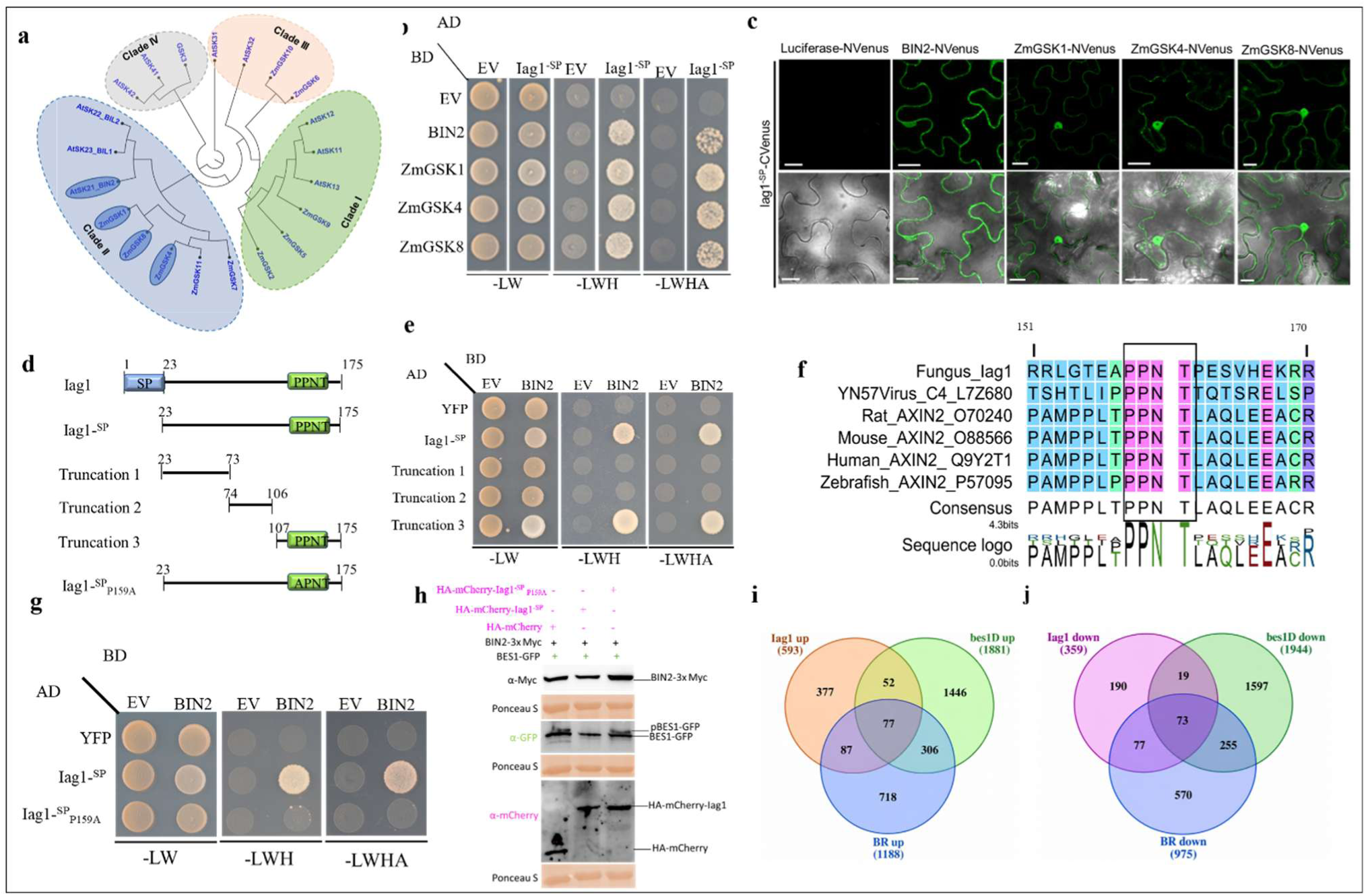
Inducer of asymmetric growth 1 (Iag1) interacts with BIN2 and maize GSK3-like kinases through a short interaction motif and attenuates BIN2-dependent signaling. **(a)** Phylogenetic analysis of Arabidopsis and maize GSK3-like kinases illustrating the relationship between BIN2 and its maize homologs. **(b)** Yeast two-hybrid assays showing interaction between Iag1 and BIN2 and its maize homologs. Growth on selective medium lacking leucine and tryptophan (-LW) selects for Y2H plasmids, whereas growth on high-stringency medium (-LWHA) indicates interaction between the indicated proteins. **(c)** Bimolecular fluorescence complementation (BiFC) assays demonstrating interaction between Iag1^-SP^ and BIN2, ZmGSK1, ZmGSK4, and ZmGSK8 in *Nicotiana benthamiana* epidermal cells. No fluorescence was detected in the negative control. Scale bar, 20 µm. **(d)** Schematic representation of Iag1 truncation constructs used to map the BIN2-interacting region and the position of the PPNT interaction motif. **(e)** Yeast two-hybrid analysis demonstrating that amino acids 107–175 of Iag1 are sufficient to mediate interaction with BIN2. **(f)** Sequence alignment of the PPNT-containing region of Iag1, Pepper yellow leaf curl virus C4, and AXIN2 proteins from Homo sapiens (human), Rattus norvegicus (rat), Mus musculus (mouse), and Danio rerio (zebrafish). These evolutionarily unrelated GSK3-interacting proteins share a PPNT sequence. **(g)** Yeast two-hybrid assays demonstrating that substitution of Proline159 abolishes interaction between Iag1 and BIN2. **(h)** Immunoblot analysis of proteins resolved on a 15% SDS-PAGE gel showing BES1-GFP phosphorylation following transient co-expression of BIN2-3×Myc with HA-mCherry, HA-mCherry-Iag1^-SP^, or the interaction-deficient mutant HA-mCherry-Iag1^-SP^_P159A_ in *Nicotiana benthamiana* leaves. **(i)** Overlap between genes upregulated following Iag1 induction and published brassinosteroid (BR)-induced and *bes1-D*-upregulated gene sets. Gene lists were compared using the complete published BR-responsive and *bes1-D* datasets (Yu et al., 2011). Statistical significance of the observed overlaps was assessed using a hypergeometric test with the complete *Arabidopsis thaliana* protein-coding gene set as the background. **(j)** Overlap between genes downregulated following Iag1 induction and published BR-repressed and *bes1-D*-downregulated gene sets (Yu et al., 2011). Statistical significance of the observed overlaps was assessed using a hypergeometric test with the complete *Arabidopsis thaliana* protein-coding gene set as the background.

To investigate the molecular basis of this interaction, we generated a series of Iag1 truncation constructs and tested their ability to bind BIN2 in Y2H assays. These experiments localized the BIN2-interacting region to amino acids 107-175 of Iag1 (Fig. 4d,e). Sequence analysis of this region revealed a PPNT motif previously identified in geminivirus C4 proteins that interact with plant GSK3-like kinases (Fig. 4f). Substitution of the central proline residue (P159A) completely abolished BIN2 binding in a yeast two-hybrid assay, demonstrating that the PPNT motif is required for the Iag1-BIN2 interaction (Fig. 4g).

BIN2 phosphorylates numerous downstream substrates involved in plant growth and development, including the transcription factor BES1. Because BES1 phosphorylation is a well-established readout of BIN2 activity, we asked whether the PPNT-dependent interaction between Iag1 and BIN2 affects this canonical output of BIN2 signaling. To test this, BIN2-3×Myc and BES1-GFP were transiently co-expressed in *N. benthamiana* leaves together with either HA-mCherry-Iag1^-SP^, the BIN2 interaction-deficient mutant HA-mCherry-Iag1^-SP^ _P159A_, or HA-mCherry as a control. Immunoblot analysis showed that co-expression of Iag1 markedly reduced the abundance of the gel-shifted phosphorylated BES1 isoform compared with the control, whereas the P159A mutant, which no longer interacts with BIN2, failed to alter BES1 phosphorylation (Fig. 4h). These results demonstrate that the PPNT motif is required for Iag1-mediated perturbation of BIN2 activity and indicate that direct interaction with BIN2 underlies the observed reduction in BES1 phosphorylation.

To determine whether the PPNT-dependent attenuation of BIN2 activity by Iag1 is reflected at the transcriptional level, we compared Iag1-responsive genes with previously published BR-responsive genes and genes differentially expressed in the constitutively active *bes1-D* gain-of-function mutant. Because reduced BIN2 activity is expected to generate transcriptional outputs resembling both BR treatment and constitutive BES1 activation, separate analyses were performed for induced and repressed genes using the complete BR-responsive and *bes1-D* gene sets from the published datasets^43^. Among the 593 genes upregulated following Iag1 induction (Fig. 3a, supplementary dataset 1), 164 were also induced by BR signaling, representing a 6.3-fold enrichment over random expectation (hypergeometric test, P = 6.0 × 10⁻⁸⁵) (Fig. 4i; Supplementary Dataset S4). Likewise, 150 of the 359 genes downregulated by Iag1 overlapped with BR-repressed genes, corresponding to an 11.6-fold enrichment (P = 2.2 × 10⁻¹²⁰) (Fig. 4j; Supplementary Dataset S5). Comparison with the *bes1-D* transcriptome revealed similarly significant concordance. Of the genes induced by Iag1, 129 were also upregulated in *bes1-D*, representing a 3.1-fold enrichment (P = 6.9 × 10⁻³²) (Fig. 4i; Supplementary Dataset S4). Likewise, 92 genes repressed by Iag1 were also downregulated in *bes1-D*, corresponding to a 3.6-fold enrichment (P = 1.6 × 10⁻²⁷) (Fig. 4j; Supplementary Dataset S5). Together, these analyses demonstrate that Iag1-responsive genes are strongly enriched among both BR-responsive and bes1-D-responsive transcriptional programs. The concordance between the Iag1 transcriptome and independently generated BR- and *bes1-D*-responsive gene sets provides independent transcriptomic evidence for attenuation of BIN2-dependent signaling.

Collectively, these findings identify BIN2 and related maize GSK3-like kinases as direct interaction partners of Iag1 and establish the PPNT motif as the molecular determinant required for this interaction. The combination of direct protein interaction, PPNT-dependent binding, reduced phosphorylation of the BIN2 substrate BES1, and the significant enrichment of BR-responsive and *bes1-D*-responsive transcriptional programs supports a model in which Iag1 attenuates BIN2-dependent signaling, thereby promoting extensive developmental reprogramming.

### Iag1 is a secreted protein of *Ustilago maydis* and is localized to the plant cytoplasm and nucleus

We next investigated whether Iag1 is secreted during fungal infection. Sequence analysis using Signal P 6.0 ^42^ predicted an N-terminal secretion signal comprising amino acids 1-23 (SP; Fig. 1c). To experimentally validate secretion, we generated *U. maydis* strains expressing C-terminal mCherry fusions of full-length Iag1 or a variant lacking the predicted signal peptide (SP), each driven by the native Iag1 promoter. Confocal microscopy of maize leaves infected with the solopathogenic strain expressing full-length Iag1-mCherry revealed fluorescence surrounding fungal hyphae, consistent with secretion of Iag1 into the biotrophic interface (Fig. 5a,b). In contrast, the SP-deletion variant (Iag1^-SP^) displayed diffuse intracellular fluorescence confined to fungal hyphae, demonstrating that the predicted signal peptide is required for secretion (Fig. 5c,d). These results establish Iag1 as a secreted *U. maydis* effector expressed during biotrophic growth.

**Figure 5.**
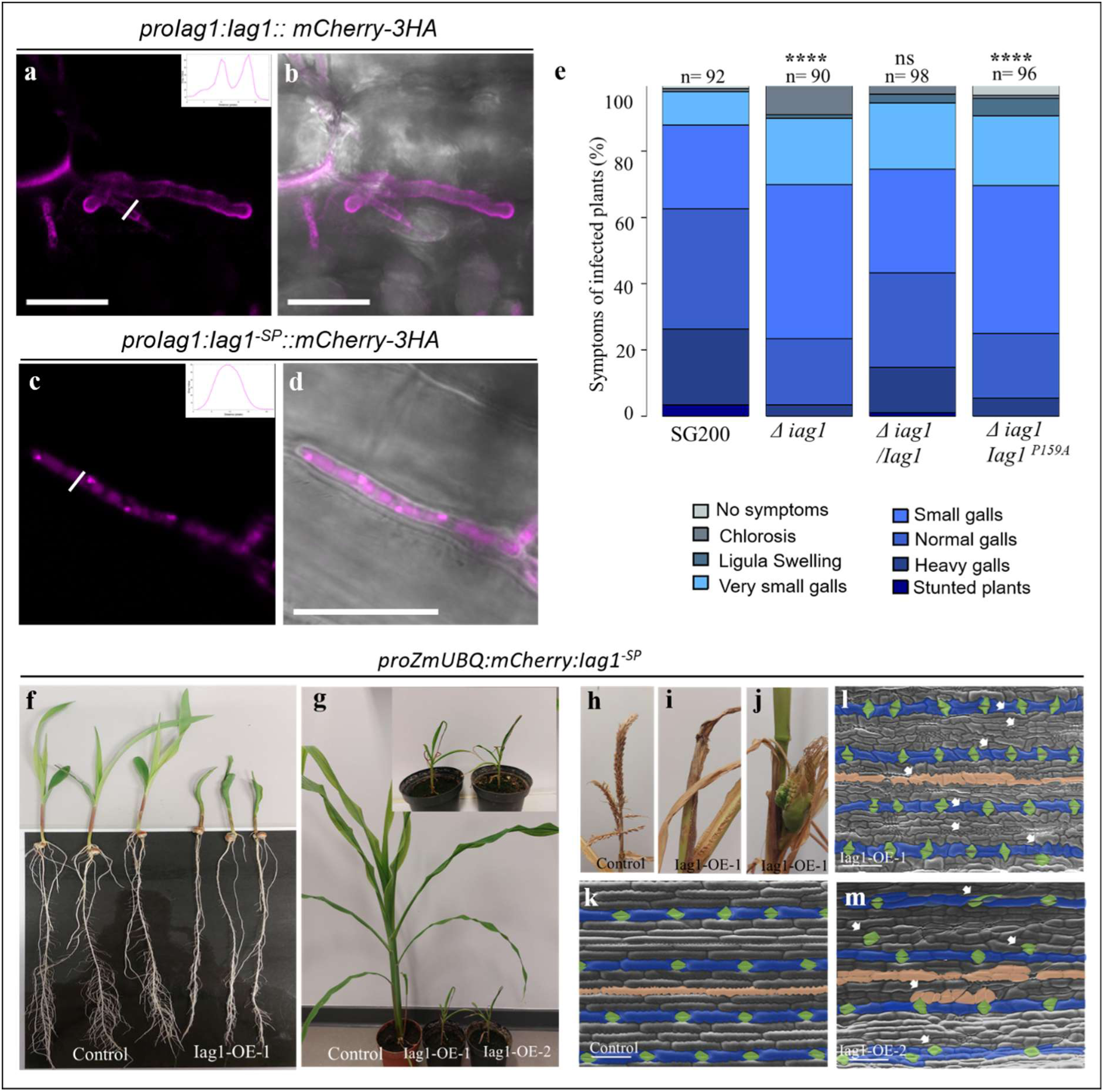
The *Ustilago maydis* effector Inducer of asymmetric growth 1 (Iag1) is secreted during infection, promotes virulence, and induces epidermal developmental reprogramming in *Zea mays*. **(a-d)** Confocal micrographs of maize leaf epidermal cells 7 days post-infection (dpi) with *U. maydis* strains expressing Iag1-mCherry-HA under the control of the native Iag1 promoter (proIag1), either with the predicted signal peptide **(a-b)** or lacking the signal peptide **(c-d)**. mCherry fluorescence surrounds fungal hyphae in the presence of the signal peptide (a,b), consistent with secretion of Iag1 into the biotrophic interface, whereas deletion of the signal peptide abolishes secretion and results in fluorescence confined to the fungal hyphae (c,d). Magenta fluorescence intensity profiles shown in the upper right corner of (a) and (c) illustrate the distribution of the mCherry signal across the hyphal diameter. **(e)** Virulence analysis of the *Δiag1* knockout mutant and complemented strains. The *U. maydis Δiag1* mutant, the wild-type Iag1 complementation strain, and the Iag1^P159A^ complementation strain were assessed for their ability to infect 8-day-old *Zea mays* (Early Golden Bantam) seedlings, using the parental SG200 strain as the control. Disease symptoms were scored 12 days post-infection (dpi). Statistical significance was determined using Fisher’s exact test (****P < 0.0001). **(f)** Representative morphology of 7-day-old transgenic maize seedlings constitutively expressing *HA-mCherry-Iag1^-SP^* or the *HA-mCherry* control under the maize ubiquitin promoter. **(g-j)** Morphological phenotypes of two independent, 8-week-old transgenic maize lines ectopically expressing *HA-mCherry-Iag1^-SP^* compared to control (*HA-mCherry*). **(k-m)** Scanning electron micrographs of the abaxial leaf epidermis from two-week-old control plants and transgenic maize expressing *HA-mCherry-Iag1^-SP^*. Blue outlines indicate stomatal lineage cells, apricot shading highlights non-stomatal epidermal cells with altered morphology, and green outlines mark guard cells. Scale bar, 200 µm.

We next investigated subcellular localization of Iag1. An N-terminal HA-mCherry fusion of Iag1^-SP^ was transiently expressed in *N. benthamiana* via *Agrobacterium tumefaciens*. Confocal microscopy revealed accumulation of *HA-mCherry-Iag1^-SP^* in both the nucleus and cytoplasm (Fig. S5a,b). A comparable localization pattern was observed in maize epidermal cells following particle bombardment, where HA-mCherry fluorescence was detected in the nucleus and cytoplasm, while co-expressed free GFP served as a marker to visualize successfully transformed plant cells and showed the expected distribution (Fig. S5c,d). Co-expression with GFP-BIN2 showed co-localization in these compartments, consistent with their physical interaction *in planta* (Fig. S5e-h). Together, these findings show that Iag1 is secreted by *U. maydis* and, when present inside plant cells, accumulates in the same subcellular compartments as its plant host target BIN2.

### Iag1 is required for full virulence of *Ustilago maydis* and induces epidermal reprogramming in maize

To determine the contribution of Iag1 to fungal virulence, we generated a CRISPR/Cas9-mediated *Δiag1* knockout mutant in the solopathogenic strain SG200 and assessed its pathogenicity in maize. Plants infected with the *Δiag1* mutant still developed galls; however, disease severity was significantly reduced compared with the parental SG200 strain, indicating that Iag1 contributes quantitatively to fungal virulence (Fig. 5e). These findings suggest that Iag1 functions as a virulence determinant that acts in concert with additional, partially redundant effectors during maize colonization, consistent with the functional redundancy characteristic of the *U. maydis* effector repertoire ^9^.

To determine whether the reduced virulence resulted specifically from loss of *Iag1* and whether the PPNT motif is required for its biological activity, the *Δiag1* mutant was complemented with either the wild-type *Iag1* allele or the interaction-deficient *Iag1^P159A^* variant. Complementation with the wild-type allele fully restored virulence to levels comparable with the parental SG200 strain, whereas expression of *Iag1^P159A^*failed to rescue the reduced disease phenotype (Fig. 5e). These results demonstrate that the PPNT motif is essential for Iag1 function during maize infection and support the conclusion that PPNT-dependent interaction with host GSK3-like kinases contributes to the full virulence of *U. maydis*.

To determine whether Iag1 is sufficient to reprogram host development in the natural host, we constitutively expressed Iag1^-SP^ in maize under the control of the maize ubiquitin promoter. Two independent transgenic lines displayed pronounced developmental abnormalities, including reduced plant stature, inward leaf curling, and complete sterility resulting from defective tassel development (Fig. 5f-j). These findings demonstrate that sustained Iag1 activity profoundly disrupts maize development, consistent with the central role of GSK3-like kinases in coordinating multiple developmental pathways.

To define the cellular basis of these phenotypes, we examined leaf epidermal organization by scanning electron microscopy. In control plants, stomata were arranged in regularly spaced longitudinal files following the characteristic one-cell spacing rule (Fig. 5k). In contrast, Iag1-expressing leaves exhibited extensive disruption of epidermal patterning, accompanied by increased cell proliferation in both stomatal and inter-stomatal cell files. This resulted in the insertion of additional pavement cells between adjacent stomata and an increased number of non-stomatal cells separating neighboring stomatal rows (Fig. 5l,m). Together, these findings demonstrate that ectopic Iag1 expression is sufficient to induce epidermal developmental reprogramming in maize, closely resembling key features of the epidermal abnormalities observed during *U. maydis* gall formation. These results support a model in which Iag1 targets maize GSK3-like kinases to reprogram host development and promote disease.

## Materials and Methods

### Molecular cloning

Coding sequences were amplified from the cDNA of *A. thaliana*, *Z. mays* (accession B73, or EGB), or *U. maydis*-infected *Z. mays* leaves by PCR, while promoter regions were amplified from the respective genomic DNA. PCR products were initially cloned into the pJET1.2/blunt vector (Thermo Fisher Scientific) and then assembled using the GreenGate cloning system (Lampropoulos et al., 2013), with the Type IIS restriction enzyme BsaI (New England Biolabs). For the generation of stable transgenic *A. thaliana* lines, coding sequences were fused to fluorescent or epitope tags (HA-mCherry or 2×Myc) and placed under the control of a β-estradiol-inducible promoter. For *U. maydis*, promoter-reporter constructs were generated by fusing the native Iag1 promoter to mCherry and integrated into the fungal genome of the saprotrophic strain SG200 ^15^ by homologous recombination as described previously ^44^. For interaction and localization assays, coding sequences of Iag1 and the indicated plant GSK3-like kinases were cloned into appropriate GreenGate-compatible destination vectors for yeast two-hybrid, bimolecular fluorescence complementation (BiFC), and plant expression as described previously in ^22,24^. All constructs were verified by Sanger sequencing prior to transformation. *Iag1* mutants were generated as described previously ^22^.

### Fungal growth conditions and infection assays

The solopathogenic *U. maydis* strain SG200 was used for the generation of the *iag1* mutant and reporter strains. Scanning electron microscopy analyses were performed using maize infected with the compatible wild-type strains FB1 and FB2 or SG200-infected maize plants. Strains were cultured in liquid YEPS medium (1% yeast extract, 2% peptone, 2% sucrose) at 28 °C with shaking at 200 rpm. For solid culture, strains were grown on potato dextrose agar (PDA) plates at 28 °C as described previously ^44^.

### Plant material and growth conditions

*A. thaliana* ecotype Columbia (Col-0) was used as the wild type for the generation of transgenic lines. Plants used for floral dipping were grown at 22 ± 2 °C under a 16 h light / 8 h dark photoperiod with a light intensity of 120-150 µmol m⁻² s⁻¹. Transformation was performed using the floral dip method as described by ^45^. Independent T1 transgenic lines expressing the effectors were screened by basta selection and western blotting, and two lines were selected for further phenotypic analysis. For phenotyping, seeds were germinated on half-strength Murashige and Skoog (½ MS) agar plates and grown for 7 days in a growth chamber (Panasonic environmental test chamber, Type: MLR-352H-PE). Seedlings were then transferred to plates supplemented with either 10 µM β-estradiol or an equivalent volume of dimethyl sulfoxide (DMSO) as a solvent control. Phenotypes were documented by non-invasive imaging of the agar plates.

*Z. mays* Early Golden Bantam was used for all infection assays, unless otherwise stated. For the infections, plants were grown in a climate chamber under controlled conditions with a 16 h light / 8 h dark photoperiod and temperatures of 28 °C (day) and 22 °C (night). Seven-day-old seedlings were infected with *U. maydis* strains, and disease symptoms were scored at 12 days post infection (dpi) according to ^15^. Statistical analysis of virulence data was performed using Fisher’s exact test in R, as described by ^46^.

*N. benthamiana* plants used for transient expression assays were grown under a 16 h light / 8 h dark photoperiod at 22-24 °C. Leaves of 4-5-week-old plants were used for *A. tumefaciens*-mediated infiltration.

### Microscopy

Confocal microscopy was performed using a Leica TCS SP8 confocal laser scanning microscope. Samples were placed on microscope slides, mounted in water, and sealed with a coverslip. mCherry fluorescence was excited at 561 nm, and emission was collected between 578 and 648 nm. For aniline blue staining, hand-cut sections of maize leaves were incubated in 0.1% (w/v) of aniline blue solution prepared in 0.07 M K₂HPO₄ buffer (pH 9.0) for 5 min in the dark. After staining, samples were washed twice with the same buffer and mounted in water. Fluorescence images were acquired using a 405 nm laser for excitation, and emission was collected between 450 and 500 nm. For propidium iodide (PI) staining, samples were incubated in a PI solution (10 µg/mL) for 5 min at room temperature, briefly washed with distilled water, and mounted in water. PI was excited at 561 nm, and emission was collected between 600 and 650 nm.

Scanning electron microscopy (SEM) was performed using an XL30 Environmental Scanning Electron Microscope (ESEM, FEI-Philips, Kassel, Germany). Samples were mounted on aluminum stubs using carbon adhesive tape. Imaging was performed under high-vacuum mode at an accelerating voltage of 10-20 kV.

### RNA Sequencing and Data Analysis

Twelve-day-old *A. thaliana* seedlings were treated with 10 µM β-estradiol for 4 h. Three independent biological replicates were collected per condition. Tissue was flash-frozen in liquid nitrogen and ground to a fine powder. Total RNA was extracted from ∼300 mg of tissue using the NEB RNA extraction kit (New England Biolabs) according to the manufacturer’s instructions, including on-column DNase treatment.

RNA-seq libraries were prepared as paired-end libraries (2 × 150 bp) and sequenced on an Illumina NovaSeq 6000 platform (Biomarker Technologies [BMK] GmbH, Münster, Germany). Raw reads were assessed for quality using FastQC (v0.12.1). Adapter sequences and low-quality bases were removed using fastp (v0.19.5). Trimmed reads were aligned to the *A. thaliana* reference genome (TAIR10 genome with Araport11 annotation) using HISAT2 (v2.1.0), retaining a single primary alignment per read (-k 1). SAM files were converted to BAM format, sorted, and indexed using samtools.

Mapped reads were assessed for each RNA-seq sample, showing 95-96% genome alignment, 24.5-34.8% duplicate reads, and an insert size distribution with a median of 273-286 bp. Gene-level read summarization was performed using featureCounts (v2.0.3; Rsubread v2.14.1), with ≥92.6% of read pairs assigned to annotated genes. Genes with a total count of 10 or fewer across all samples were removed prior to downstream analysis.

Differential gene expression analysis was performed in R using DESeq2 (v1.40.1). The experimental condition (Iag1^-SP^ versus control) was specified as the factor in the statistical model. Genes with an adjusted P-value ≤ 0.05 and |log2 fold change| ≥ 1 were considered differentially expressed. Volcano plots and heatmaps were generated in R using ggplot2 and pheatmap, respectively, after a variance-stabilizing transformation of count data, with hierarchical clustering based on Euclidean distance.

Gene Ontology (GO) and pathway enrichment analyses were performed using ShinyGO (v0.85.1; https://bioinformatics.sdstate.edu/go/). Separate analyses were conducted for upregulated and downregulated gene sets. Enrichment was assessed using a hypergeometric test with false discovery rate (FDR) correction, and terms with adjusted p ≤ 0.05 were considered significant.

To assess whether Iag1-regulated genes significantly overlapped previously published BR-responsive and bes1-D-responsive gene sets ^43^, overlap enrichment was evaluated using a hypergeometric test. The background population consisted of all annotated Arabidopsis protein-coding genes (Araport11).

### Co-immunoprecipitation and mass spectrometry

Seven-day-old *A. thaliana* seedlings were transferred to ^1^_2_ MS plates containing 10 µM β-estradiol for five days. 0.4 g of seedlings were harvested, shock-frozen in liquid nitrogen, and ground using a mortar and pestle. Two independent biological replicates were collected and total proteins were extracted as described previously ^47^. Total protein extracts were immunopurified using anti-c-Myc antibody coupled to magnetic beads (MACS® Technology, Miltenyi). Proteins were digested on-column with trypsin, and the resulting peptides were subjected to LC-MS/MS analysis in a single run ^48^.

The resulting tryptic peptide mixture was desalted before LC-MS/MS analysis on a C18 ZipTip (Omix C18 100 µL tips, Varian). The purified peptide mixture was analyzed by LC-MS/MS using a nanoflow RP-HPLC system (LC program: linear gradient of 3-40% solvent B over 100 min; solvent A: 0.1% formic acid in water; solvent B: 0.1% formic acid in acetonitrile) on-line coupled to an Orbitrap mass spectrometer (Orbitrap Fusion Lumos, Thermo Fisher Scientific) operating in positive ion mode.

The 20 most abundant multiply charged precursor ions from each MS survey scan were selected for MS/MS analysis. Both MS and MS/MS spectra were acquired in the Orbitrap, and peptide fragmentation was performed using higher-energy collisional dissociation (HCD). Raw data were processed into peak lists using Proteome Discoverer (v1.4) and searched against the *A. thaliana* protein database (UniProt) using the Protein Prospector search engine (v5.15.1). The following search parameters were applied: enzyme specificity: trypsin with a maximum of 2 missed cleavages; mass tolerances: 5 ppm for precursor ions and 10 ppm for fragment ions (both monoisotopic); fixed modification: carbamidomethylation of cysteine residues; variable modifications: protein N-terminal acetylation, methionine oxidation, and cyclization of N-terminal glutamine residues, allowing a maximum of 2 variable modifications per peptide. Acceptance criteria were as follows: minimum protein and peptide scores of 22 and 15, respectively; maximum E-values of 0.01 for proteins and 0.05 for peptides. Spectral counting was used to estimate relative protein abundance in the no-antibody and c-Myc-GFP negative controls and in the anti-c-Myc immunopurified samples.

### Generation of transgenic *Zea mays* lines

The *HA-mCherry-Iag1^-SP^* coding sequence was cloned into the binary vector pCAMBIA3300 under the control of maize ubiquitin1 promoter and the nopaline synthase (NOS) terminator. An HA-mCherry construct was used as a control, resulting in pUbi-HA-mCherry. The vectors carry the bar gene for selection of transgenic events. Transgenic plants of the maize inbred line B104 were generated via *Agrobacterium tumefaciens*-mediated transformation of immature embryos using *HA-mCherry-Iag1^-SP^* or the control construct *pUbi-HA-mCherry*, by using the established protocol ^49^. T0 plants were grown to maturity in a greenhouse, and the subsequent generation was used for molecular and phenotypic analyses.

## Discussion

Pathogen-induced gall formation in plants represents a striking example of host developmental reprogramming; however, the molecular mechanisms by which pathogens manipulate host growth remain incompletely understood. Previous studies demonstrated that gall formation in *U. maydis* predominantly involves mesophyll- and bundle sheath-derived cells ^30^; much less was known about how epidermal tissues respond to infection. Here, we show that the epidermis also undergoes extensive developmental reprogramming, characterized by altered cell division orientation, disruption of stomatal patterning, and profound changes in cell morphology. Our findings therefore identify epidermal developmental reprogramming as an additional and previously underappreciated component of gall development. We further identify the secreted *U. maydis* effector Iag1 as a regulator of this process and show that it promotes gallogenic growth by targeting conserved host GSK3-like kinases. Together, these findings establish a direct mechanistic link between a fungal effector and a central host developmental signaling hub, demonstrating how manipulation of BIN2 and related kinases can reprogram cell fate, growth behavior, and tissue organization during gall formation.

BIN2 is increasingly recognized as a central signaling hub instead of simply a negative regulator of BR signaling. In addition to phosphorylating the transcription factors BES1 and BZR1, BIN2 regulates numerous substrates involved in auxin signaling, stomatal development, cell division, and cell fate specification ^33,50^. Our biochemical and transcriptomic analyses indicate that Iag1 perturbs GSK3-like kinase activity through direct physical interaction. Reduced phosphorylation of the canonical BIN2 substrate BES1 provides one biochemical readout of this perturbation, while the strong enrichment of BR- and *bes1-D*-responsive genes provides independent evidence for altered BIN2-dependent signaling. Importantly, the developmental phenotypes induced by Iag1 extend well beyond those typically associated with BR treatment or mutations affecting BR biosynthesis or perception. Iag1 expression induces extensive epidermal reprogramming, including altered division orientation, disruption of stomatal patterning, and widespread changes in cell morphology. These broader developmental effects are likely to reflect the central role of BIN2 and related GSK3-like kinases in coordinating multiple signaling pathways rather than BR signaling alone. Consistent with this interpretation, the Iag1 transcriptome revealed extensive changes in auxin-responsive transcription in addition to its strong BR- and *bes1-D*-like signature. Together, these findings suggest that Iag1 exploits GSK3-like kinases as developmental signaling hubs instead of simply activating canonical BR signaling. By perturbing multiple GSK3-like kinase-dependent outputs, including BR and auxin signaling and potentially additional developmental pathways, Iag1 may unlock a degree of developmental plasticity that cannot be explained by altered BR signaling alone. This broader rewiring of GSK3-like kinase-dependent signaling provides a mechanistic framework for the extensive cellular reprogramming associated with gall formation.

A notable finding of our study is the identification of a PPNT motif in Iag1 that is essential for BIN2 binding, as substitution of a single proline residue abolished both interaction with BIN2 and the virulence factor activity of Iag1 during infection. The same motif is present in geminiviral C4 effectors that also target GSK3-like kinases (Fig. 4f) ^40^, suggesting that evolutionarily unrelated plant pathogens have converged on a common molecular strategy to manipulate this central signaling hub. These observations identify the PPNT motif as a candidate GSK3 interaction motif that may be shared among diverse pathogen effectors.

Intriguingly, we also identified a PPNT sequence within the metazoan scaffold protein AXIN2, a core regulator of GSK3 in the β-catenin destruction complex ^51^. This sequence is located within a predicted intrinsically disordered region of AXIN2, consistent with the enrichment of short linear motifs (SLiMs) in flexible protein regions that mediate transient protein-protein interactions ^52^. Although the functional significance of this sequence in AXIN2 has not been experimentally established, its presence raises the possibility that PPNT-like motifs represent a more general mechanism for engaging GSK3-family kinases. More broadly, our findings suggest that short linear motifs embedded within intrinsically disordered regions provide an evolutionarily flexible means by which both endogenous regulatory proteins and pathogen effectors can access conserved kinase interaction surfaces. Testing whether these motifs recognize analogous structural interfaces on plant and metazoan GSK3 proteins will require future biochemical and structural studies.

Targeting central negative regulators has emerged as a recurring strategy by which microbial effectors reprogram host signaling. Instead of directly activating developmental pathways, effectors frequently manipulate endogenous signaling hubs that normally constrain transcriptional and developmental responses. In *U. maydis*, this strategy is exemplified by Nkd1, Jsi1 and members of the Tip family, which activate transcriptional programs by interfering with the TOPLESS co-repressor complex ^10,18,21,22,24,25^, as well as Hap1, which promotes susceptibility through inhibition of the metabolic regulator SnRK1 ^53^. Our findings extend this emerging paradigm by identifying BIN2 as an additional regulatory hub targeted during infection. Unlike previously described effectors that primarily modulate transcriptional repression or metabolic signaling, Iag1 targets a multifunctional developmental kinase that integrates hormone signaling, cell fate specification and tissue patterning. This highlights BIN2 as a particularly effective point of intervention through which a single effector can simultaneously influence multiple developmental programs.

Functional analyses in the natural host establish Iag1 as a quantitatively important virulence determinant. Deletion of *Iag1* significantly reduced disease severity, whereas complementation with the wild-type allele restored virulence, confirming that the observed phenotype results from loss of Iag1. In contrast, complementation with the BIN2 interaction-deficient *Iag1^P159A^*allele failed to rescue virulence, demonstrating that the PPNT motif is required for Iag1 function during infection. Although loss of Iag1 attenuated disease, tumor formation was not abolished, indicating that developmental reprogramming is controlled by multiple partially redundant effectors. This conclusion is consistent with previous studies of See1, which promotes host cell proliferation by targeting the Suppressor of G2 allele of SKP1 (SGT1), and the recently described effector Tip4, which promotes proliferative growth by activating a lateral root induction-like developmental program ^28^. Interestingly, Tip4-induced developmental phenotypes have so far been observed primarily in Arabidopsis roots, whereas stable transgenic maize lines expressing Tip4 could not be regenerated, precluding direct assessment of its developmental activity in maize. By contrast, Iag1 induces pronounced developmental defects in aerial plant tissues including maize, suggesting that distinct effectors converge on host developmental reprogramming through complementary mechanisms operating in different developmental contexts.

Together, our findings identify GSK3-like kinases as previously unrecognized targets *of U. maydis* effectors and demonstrate that perturbation of GSK3-like kinase signaling is sufficient to trigger extensive developmental and transcriptional reprogramming. Instead of simply activating BR signaling, Iag1 exploits BIN2 as a central developmental signaling hub to coordinately rewire multiple downstream pathways, providing a mechanistic explanation for the remarkable developmental plasticity underlying gall formation. More generally, our work supports a model in which pathogen effectors manipulate highly connected regulatory hubs rather than individual signaling pathways to efficiently reprogram host development and promote disease. The identification of a PPNT interaction motif further suggests that short linear motifs provide an evolutionarily flexible mechanism by which diverse pathogens independently converge on GSK3-family kinases to reprogram host development.

## Author contributions

M.K. and A.D. conceived the study. M.K. designed and performed most experiments and analyzed the data. A.P.Z. performed co-immunoprecipitation and mass spectrometry analyses to identify interaction partners. P.B. contributed to the generation and phenotypic analysis of transgenic Arabidopsis lines. G.L. and A.Z. performed RNA-seq analysis and interpretation. H. L. and B.Y. designed and constructed *Iag1* expression and control vectors, and generated transgenic maize lines. M.K. and A.D. supervised the study and secured funding. M.K. and A.D. wrote the manuscript with input from all authors.

## Acknowledgements

We thank Dr. Evan John for the critical reading of the manuscript and helpful discussions. We are grateful to Knut Wichterich for assistance with scanning electron microscopy. This work was supported by the Deutsche Forschungsgemeinschaft (DFG, German Research Foundation) under Germany’s Excellence Strategy EXC-2070-390732324 and DFG grants (435107456, 536218570, 520490591, 557819931).

