## Supplementary data for "A fungal effector hijacks conserved plant GSK3-like kinases to reprogram host cell identity"

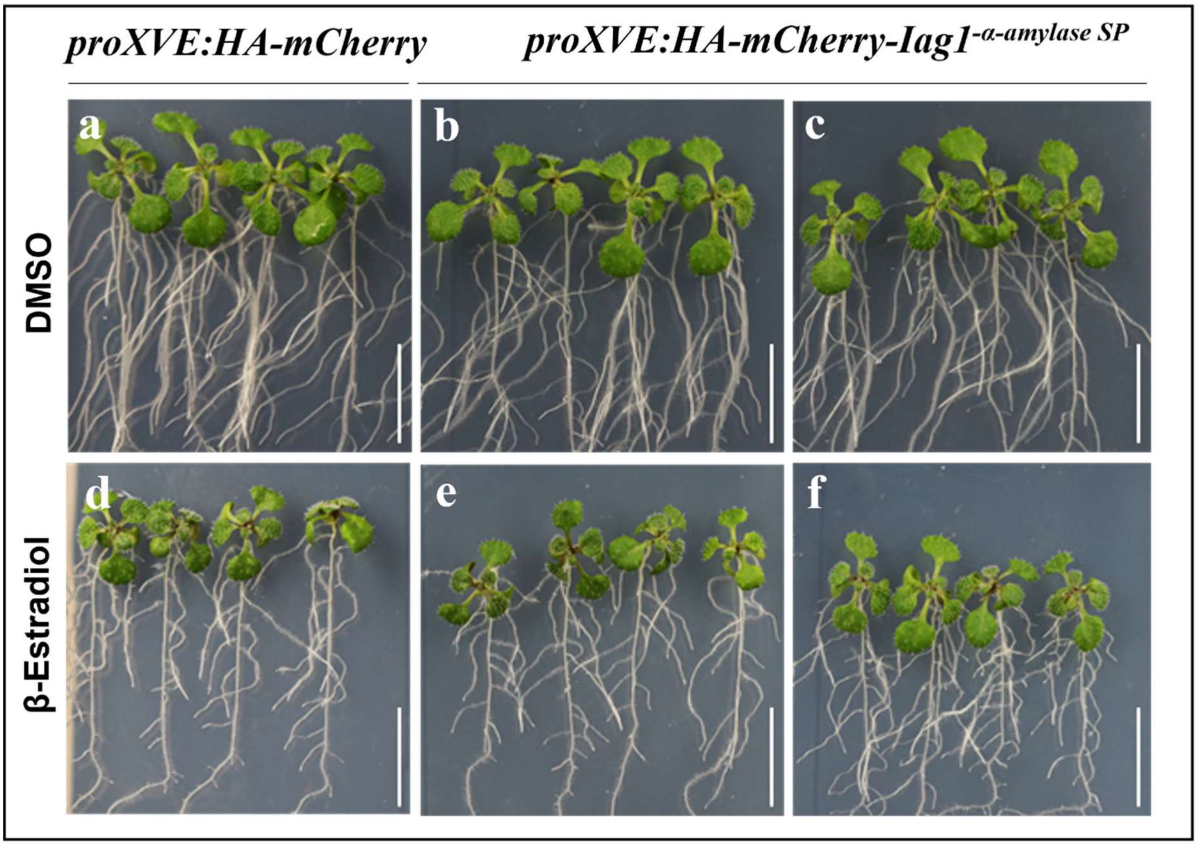


**Supplementary Figure 1. Expression of plant-secreted fungal effector Iag1 does not induce developmental phenotypes in *Arabidopsis thaliana*.** Morphological phenotypes of two independent transgenic *A. thaliana* lines expressing Iag1 fused to a plant signal peptide for secretion into the apoplast compared with HA-mCherry-expressing control plants. Images were acquired 7 days after induction with 10 μM β-estradiol. No detectable developmental abnormalities were observed following induction of the secreted Iag1 construct.

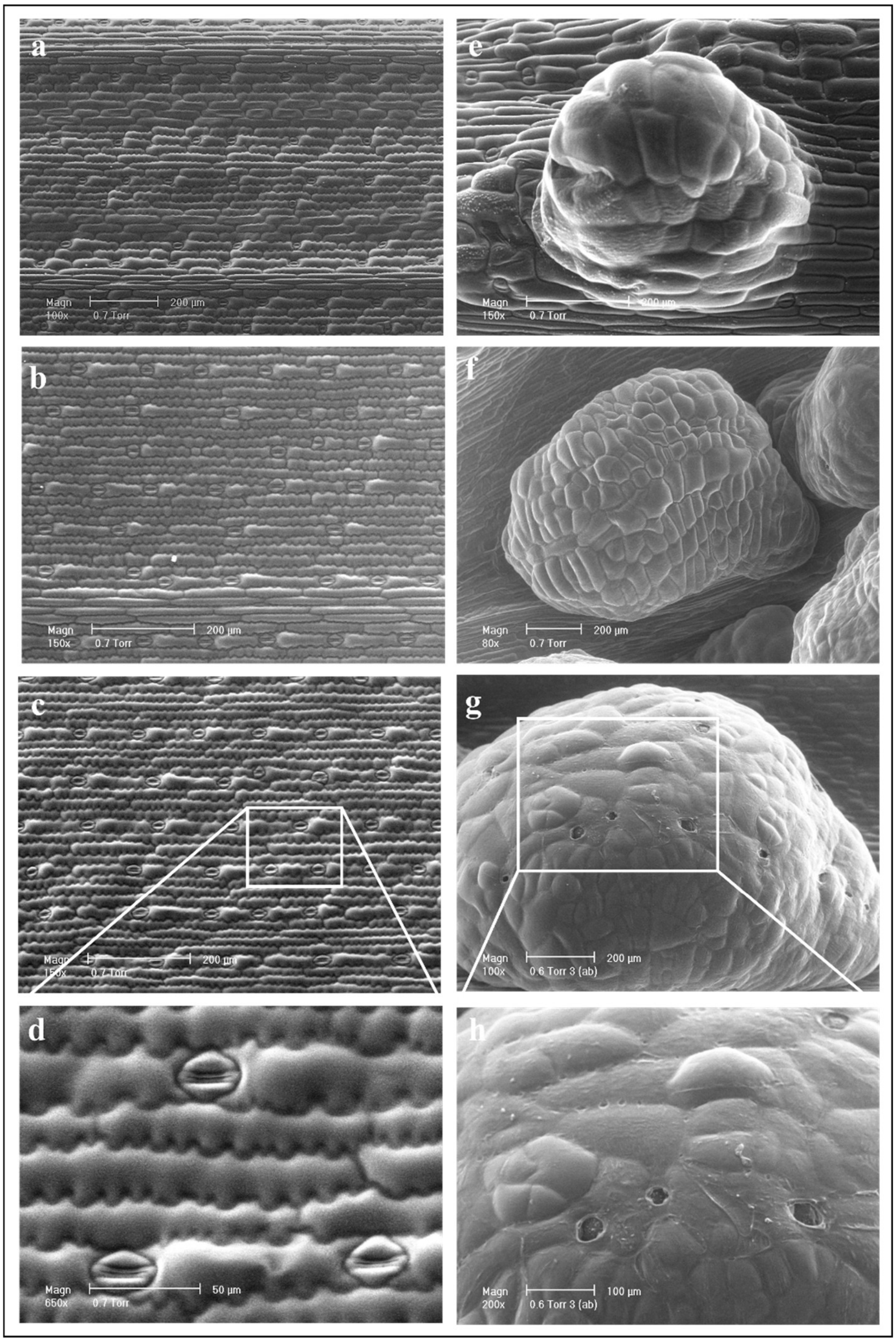


**Supplementary Figure 2.** ***Ustilago maydis*** **infection disrupts epidermal cell morphology and stomatal patterning in maize leaves. (a-d)** Scanning electron micrographs of the abaxial leaf epidermis from mock-inoculated maize leaves. **(e-h)** Scanning electron micrographs of the abaxial leaf epidermis from *U. maydis*-infected maize leaves. Images were acquired 7 days post-inoculation (dpi). **(d)** Higher-magnification view of a representative stomata in a mock-treated leaf. **(h)** Higher-magnification view of a representative pore-like opening remaining within gall tissue following *U. maydis* infection.

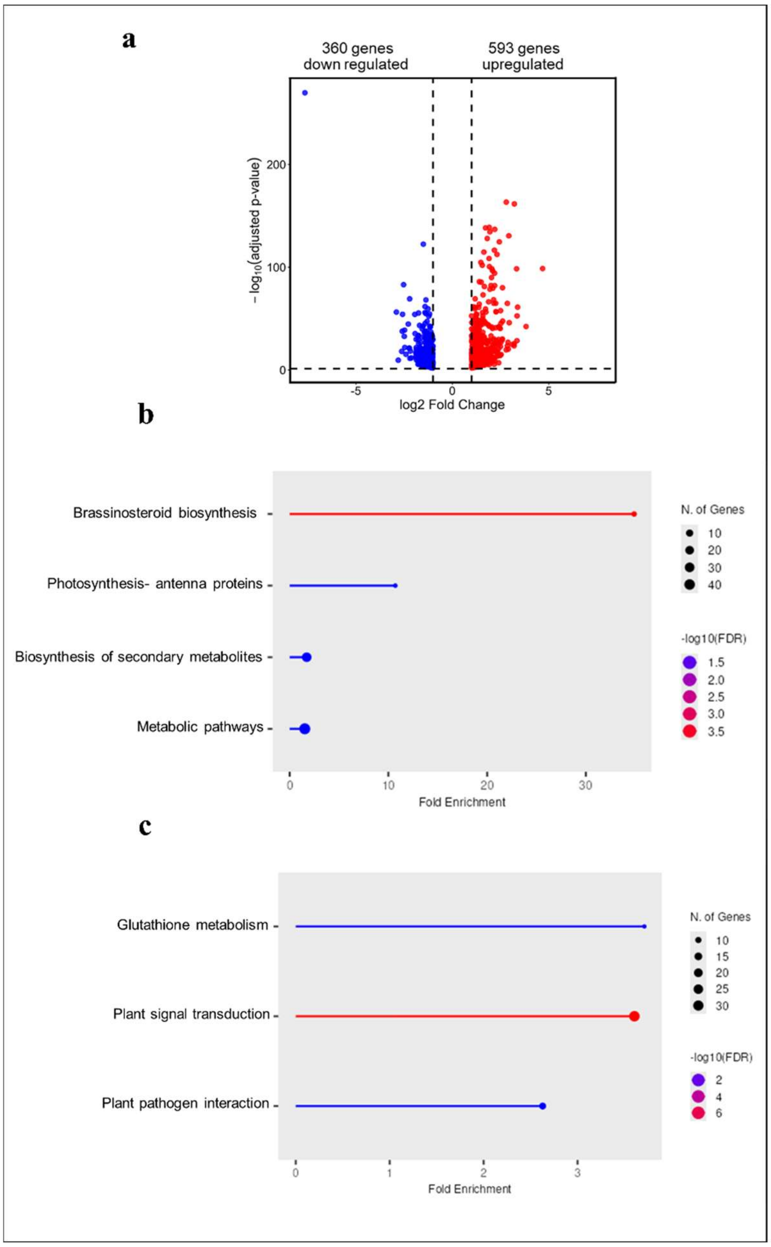


**Supplementary Figure 3. Transcriptomic analysis of *Arabidopsis thaliana* expressing Iag1. (a)** Volcano plot showing differentially expressed genes in *A. thaliana* seedlings expressing HA-mCherry-Iag1$^{-SP}$ compared with identically treated HA-mCherry control plants. Genes with an adjusted $P \leq 0.05$ and $|\log_2$ fold change$| \geq 1$ are highlighted as significantly upregulated or downregulated. **(b)** Gene Ontology (GO) enrichment analysis of significantly downregulated genes. **(c)** Gene Ontology (GO) enrichment analysis of significantly upregulated genes. Dot plots show enriched GO terms associated with biological processes. The x-axis indicates fold enrichment. Dot size represents the number of genes associated with each GO term, and color indicates statistical significance ($-\log_{10}$ FDR), with red denoting greater significance.

**a**

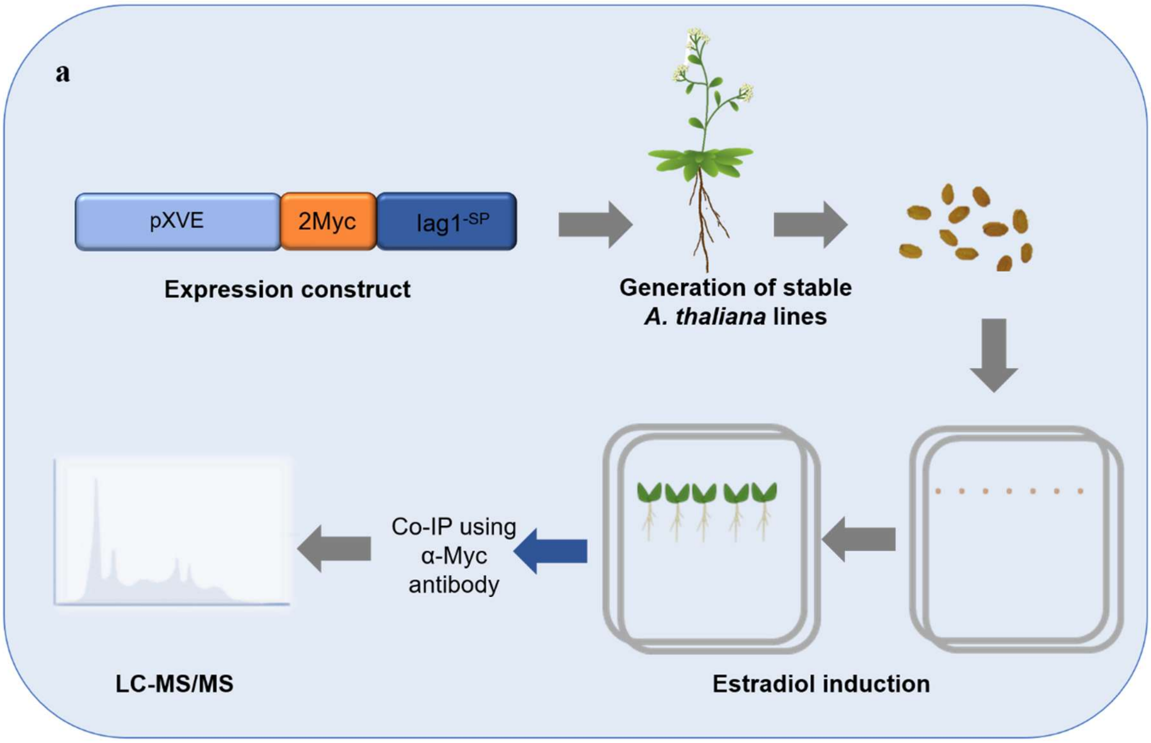


**b**

| No. | Names of putative Interactor | ATG numbers |
|---|---|---|
| 1 | CYSTATIN1 | AT5g12140 |
| 2 | SAP18 | At2g45640 |
| 3 | GLYCINE-RICH PROTEIN 2b | AT2g21060 |
| 4 | BIN2 | AT4G18710 |

**c**

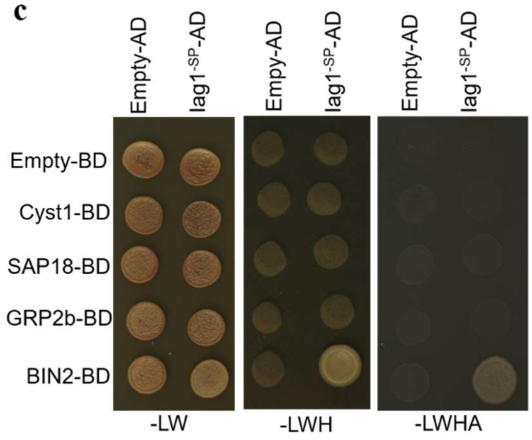


**Supplementary Figure 4: Proteomic identification and validation of protein interaction partners. (a)** Experimental workflow for interaction mapping. *Arabidopsis thaliana* lines expressing a 2xMyc-tagged protein under a β-estradiol-inducible promoter were generated. Seven-day-old seedlings were transferred to 10 µM β-estradiol-containing medium for 5 days to induce protein expression before co-immunoprecipitation (co-IP). Immunoprecipitated complexes were analysed by mass spectrometry to identify associated proteins. **(b)** Candidate interactors identified by mass spectrometry analysis. **(c)** Direct interaction validation by pairwise yeast two-hybrid assays. Growth on selective medium lacking leucine and tryptophan (-LW) indicates transformation efficiency, whereas growth on high-stringency medium (-LWHA) indicates protein-protein interaction.

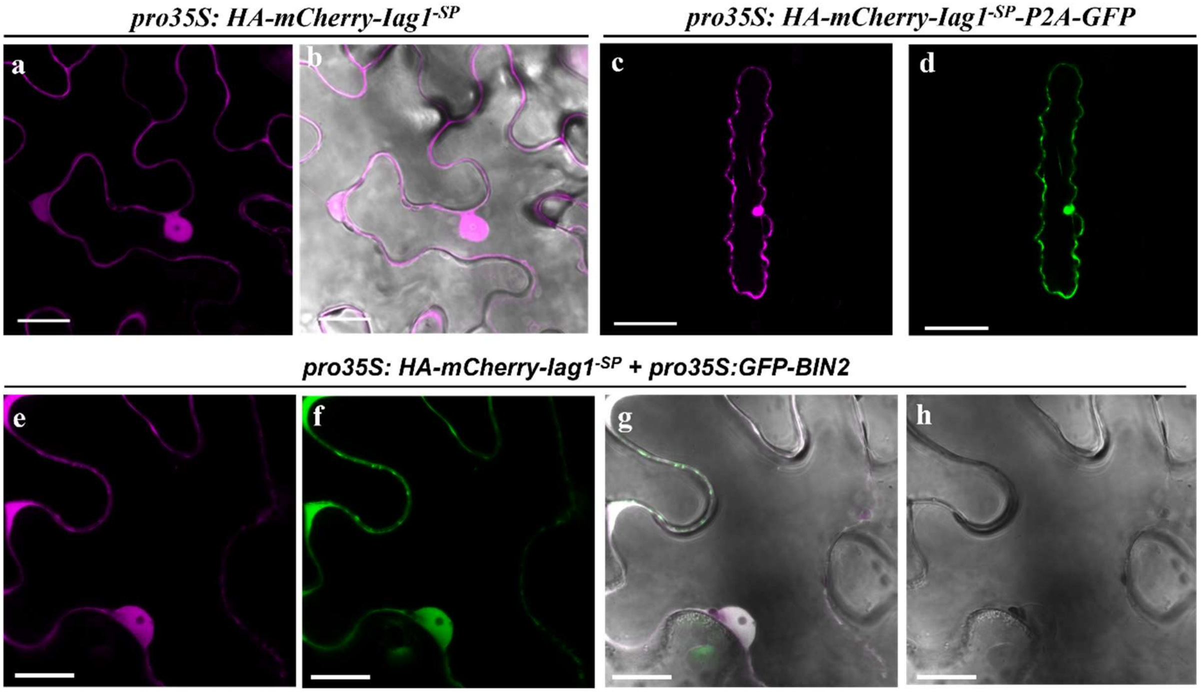


**Supplementary Figure 5. Subcellular localization of Iag1 in plant cells. (a,b)** Transient expression of HA-mCherry-Iag1$^{-SP}$ in *Nicotiana benthamiana* leaf epidermal cells under the control of the CaMV 35S promoter (pro35S), showing localization to both the cytoplasm and nucleus. **(c,d)** Subcellular localization of HA-mCherry-Iag1$^{-SP}$ following transient expression in maize leaf epidermal cells by particle bombardment. Free GFP was co-expressed via a P2A ribosomal skipping peptide as a transformation marker to identify transformed cells. **(e-h)** Co-localization of GFP-BIN2 and HA-mCherry-Iag1$^{-SP}$ following transient co-expression in *Nicotiana benthamiana* leaf epidermal cells. Scale bars, 50 µm.
